# The C-terminal tail of CSNAP attenuates the CSN complex

**DOI:** 10.1101/2022.07.18.500399

**Authors:** Maria G. Füzesi-Levi, Gili Ben-Nissan, Dina Listov, Zvi Hayouka, Sarel Fleishman, Michal Sharon

## Abstract

Protein degradation is one of the essential mechanisms that enables reshaping of the proteome landscape in response to various stimuli. The largest E3 ubiquitin ligase family that targets proteins to degradation by catalyzing ubiquitnation is the cullin-RING ligases (CRL). Many of the proteins that are regulated by CRLs are central to tumorigenesis and tumour progression, and dysregulation of the CRL family is frequently associated with cancer. The CRL family comprises ∼300 complexes all of which are regulated by the COP9 signalosome complex (CSN). Therefore, the CSN is considered an attractive target for therapeutic intervention. Research efforts for targeted CSN inhibition have been directed towards inhibition of the complex enzymatic subunit, CSN5. Here, we have taken a fresh approach focusing on CSNAP, the smallest CSN subunit. Our results show that the C-terminal region of CSNAP is tightly packed within the CSN complex, in a groove formed by CSN3 and CSN8. We show that a 16 amino acid C-terminal peptide, derived from this CSN interacting region, can displace the endogenous CSNAP subunit from the complex. This, in turn, leads to a CSNAP null phenotype that attenuates CSN activity and consequently CRLs function. Overall, our findings emphasize the potential of a CSNAP-based peptide for CSN inhibition as a new therapeutic avenue.

## Introduction

All cells depend on a balanced and functional proteome, i.e. proteostasis, which enables adaptation to external and internal perturbations^1^. This highly sophisticated interconnected system involves protein synthesis, folding and degradation. The functionality of the proteostasis system requires high specificity as different cell types that are structurally and functionally diverse exhibit distinct proteomes^2^. In the case of protein degradation, specificity is mainly achieved by E3 ligases that ubiquitinate distinct proteins, resulting in their degradation^3^.

Eukaryotic cells express hundreds of ubiquitin E3 ligases, which can operate in different cellular contexts, respond to numerous cellular signals, and process diverse protein substrates^4^. One of the largest E3 ubiquitin ligase families that account for nearly half of all E3 ligases and is responsible for ubiquitination of 20% of the proteins degraded by the 26S proteasome, comprises cullin-RING ligases (CRLs)^5^. CRL are modular protein complexes that are assembled around a central cullin scaffold, which is associated with a specific substrate receptor, adaptor protein, and a RING protein that recruits the E2 enzyme^6^. The nine human cullins and their interchangeable set of substrate specificity factors and adaptors generate about ∼300 different configurations of CRL complexes. These complexes coordinate the levels of numerous proteins affecting every facet of eukaryotic cellular processes, such as cell-cycle, cellular proliferation, hypoxia signaling, reactive oxygen species clearance and DNA repair. Remarkably, in spite of the great diversity of CRLs in terms of composition and function all of them are regulated by the COP9 signalosome complex (CSN)^7^.

The 9-subunit CSN complex regulates CRLs by a dual inhibitory mechanism involving catalytic and non-catalytic functions. The first involves the catalytic subunit CSN5 that enzymatically deconjugates the ubiquitin-like protein Nedd8 from the cullin subunit (deneddylation)^8^. The deneddylation of CRLs promotes new substrate binding and prevents autoubiquitination, which may lead to degradation of the substrate recognition subunits of the CRL. The second mechanism is mediated through physical binding to CRLs, sterically precluding interactions with E2 enzymes and ubiquitination of substrates^9,10^. Together, these two mechanisms control the dynamic assembly and disassembly cycles of CRLs that enable adaptation of the complex configuration to the cellular needs.

As a direct regulator of CRLs, the CSN is considered an attractive drug target. Major efforts towards this direction were focused on CSN5, the catalytic subunit of the complex. For example, methods for silencing CSN5 gene expression were established^15-17^. Moreover, various screening assays were developed for identifying compounds that specifically decrease the deneddylation activity of CSN5^11-14^. As a result, compounds that inhibit the deneddylase activity of CSN5 and in turn inhibit tumor cell growth have been identified as, doxycycline, an inexpensive, commonly used and well-tolerated antimicrobial agent^18^, and the zinc binding inhibitor CSN5i-3^18-22^. So far, however, this has not resulted in CSN5 inhibitors that are ready for clinical use.

Our discovery of CSNAP, the ninth integral subunit of the CSN complex, led us to propose this subunit as a new therapeutic avenue^23,24^. CSNAP, which exists in one-to-one stoichiometry with the other CSN subunits, consists of only 57 amino acids (molecular weight: 6.2 kDa) (Fig. 1A). Its structural characteristics indicate mostly random-coil with transiently formed extended structure in the C-terminus^25^. Moreover, cross-linking mass spectrometry (MS) indicates that CSNAP binds the CSN in a cleft formed by CSN1, CSN3, and CSN8, resulting in local subunit reorientations^26^ and it was shown that the C-terminal region is essential for its integration into the complex^24^. In addition, data indicates that CSNAP attenuates CSN interactions with CRL, affecting the dynamic plasticity of CRL configuration^23^. Consequently, the absence of CSNAP (CSN^ΔCSNAP^) alters cell cycle progression and reduces cellular viability. Taking together these observations, we hypothesized that by preventing CSNAP integration within the CSN complex, i.e. mimicking the CSN^ΔCSNAP^ characteristics the functionality of the complex could be inhibited, impairing cell cycle progression, proliferation and the adaptive response to oncogenic stress conditions^23^.

**Figure 1.**
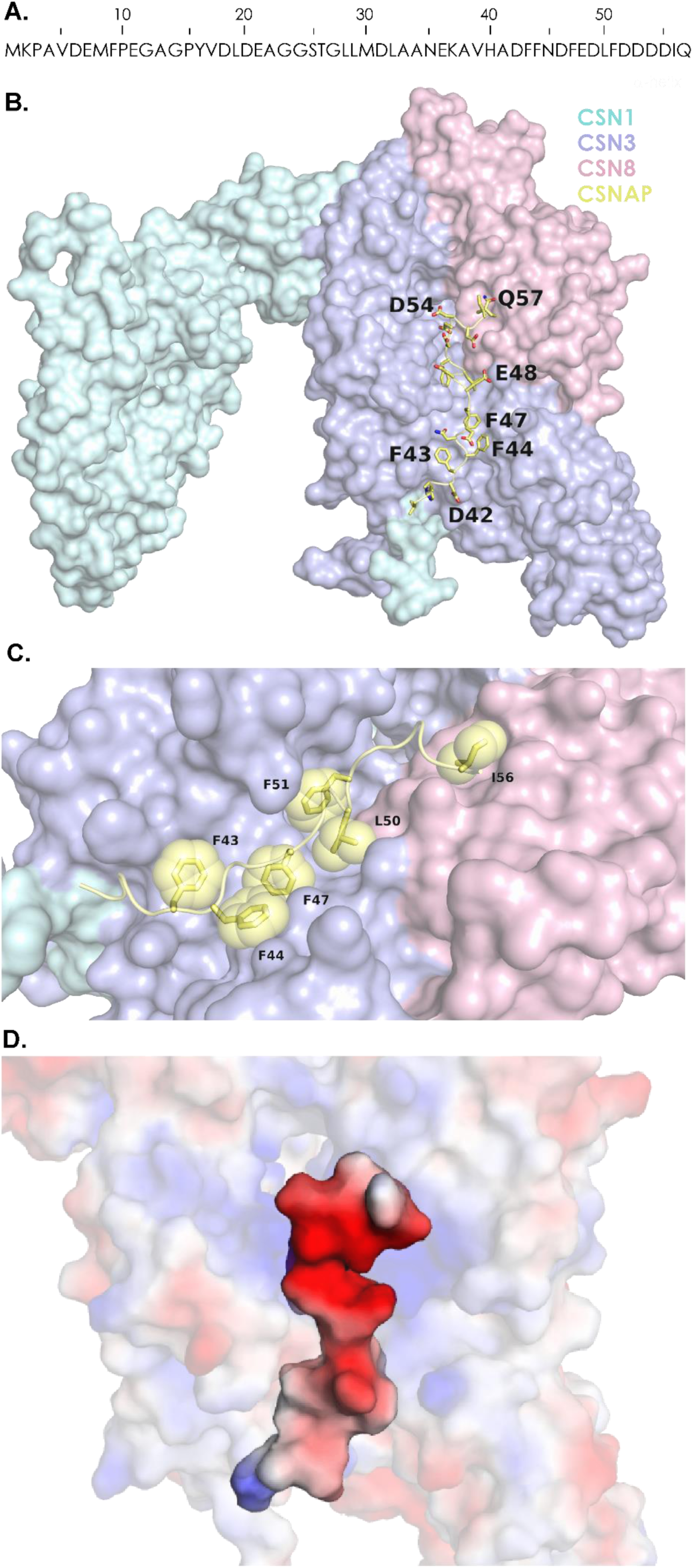
The CSNAP C-terminus binds within a groove formed by CSN3 and CSN8. A) The sequence of CSNAP. B) AlphaFold prediction of the CSN1, CSN3, CSN8 and the C-terminal region of CSNAP structure. The surfaces of CSN1, CSN3 and CSN8 are colored in cyan, purple and pink respectively. The C-terminal region of C-CSNAP, which displayed high confidence is represented by sticks colored in yellow. C) Close-up of CSNAP’s binding region. The hydrophobic and aromatic residues of CSNAP (shown as yellow spheres) are largely buried within the groove formed by CSN3 and CSN8. D) Electrostatic potential distribution maps of CSN3, CSN8 and CSNAP colored by charge (red negative and blue positive). Maps for CSN3/CSN8 and CSNAP were generated separately. The image emphasize the negative charge of CSNAP and the complementary positive potential of the CSN3/CSN8 groove.

Here, we show that the cellular presence of a C-terminal CSNAP (C-CSNAP) peptide, prevents the incorporation of the endogenous CSNAP subunit into the complex. Preventing CSNAP association leads to reduced cellular proliferation, as occurs in the CSNAP null phenotype harboring attenuated CSN function^24^. Thus, the results open up a new avenue for CSN inhibition. Since an experimentally determined structure for CSNAP in complex with CSN is not yet known, our strategy was based on generating an AlphaFold2 model and challenging it with partial structural and cross-linking data^27^. Based on the predicted structure we selected the C-CSNAP peptide and screened for its optimal properties using peptide array analysis. This approach unravels various properties required from the inhibitory peptide and further supports the predicted CSNAP-bound CSN structure. Overall, our results suggest that preventing CSNAP integration within the CSN complex by an inhibitory peptide opens a new therapeutic avenue for CSN inhibition.

## Results

### The C-terminal tail of CSNAP forms the main interaction region with the CSN complex

In solution, CSNAP is an intrinsically unstructured protein, with a slight tendency to form an extended structure in its C-terminus^25^. Within the context of the intact complex, however, the conformation of this subunit is unknown. To assess the CSNAP bound state, we predicted its structure using AlphaFold2 (AF) which has recently demonstrated atomic-level accuracy in *ab initio* prediction of protein complexes^28,29^. Considering a recent cross-linking mass spectrometry (MS) study that showed that CSNAP is positioned in a cleft formed among CSN1, CSN3, and CSN8^26^, we generated a model of these three subunits together with CSNAP . The predicted four subunit structure was superimposed onto the X-ray crystallographic structure of the CSN^ΔCSNAP^ structure (PDB 4D10)^27^, showing remarkable similarity with an RMSD of 1.02 and 0.62 Å for CSN1 and CSN3/CSN8, respectively, and 2.3 Å for the three subunits together. We further evaluated prediction quality based on the AF confidence parameters of ipTM, which is a measure of overall predicted model accuracy weighted more heavily at the oligomeric interfaces, and predicted aligned error (PAE) which describes the confidence in the relative orientation of the monomers. The predicted four subunit structure displayed a convergent interaction interface with ipTM scores of 0.83 and favorable PAE values along CSN1, CSN3, CSN8 and the CSNAP C-terminal interfaces (Fig. 1B and Supplementary Fig. 1). Judging by the low per-residue estimate of AF confidence (plDTT scores) of the CSNAP’s N-terminal region, we estimate that this region is unstructured, and we have omitted it from the structure. This is consistent with our previous results indicating that deletion of the N-terminal region of CSNAP, involving the first 20 amino acids, has no effect on CSNAP integration into the CSN complex^24^. Additional support for the importance of the C-terminal region for forming the CSNAP-bound CSN complex, comes from the cross-linking MS study in which 14 out of the 16 CSNAP containing cross-links were positioned at the C-terminal region of CSNAP^26^.

The model further supports our previous finding that Phe44 and Phe51 are necessary mediators of CSNAP interaction with the CSN^24^ (Fig 1C). These aromatic residues are buried within the CSN3 *α*-helix bundle forming the CSNAP/CSN interface. Moreover, as suggested^24^, an amphipathic configuration is formed, in which the aromatic residues face the core of the complex and the negatively charged residues are exposed to the solvent (Fig. 1D). We then mapped the previously determined cross-linking constraints^26^ of C-CSNAP on the generated model. Cross-links involving the C-terminus of CSN1 (positions 472 and 477) were not considered, due to the multiple degrees of freedom of this region, whereas the linkages involving CSN3 satisfied all 8 constraints. The distance between CSN3 Glu133 and CSNAP Asp42, Asp46, Asp48 and Asp49 was below 25 Å, and likewise, the distance between CSN3 Glu333 and CSNAP Asp52, Asp53, Asp54, Asp55 was below 20 Å. Taken together, the results indicate that C-CSNAP forms the main binding interface with the CSN complex, leading us to suggest that a synthetic C-terminal peptide that will accommodate this region, may preclude the integration of CSNAP into the complex, thereby generating the CSN^ΔCSNAP^ phenotype wherein the complex function is inhibited.

### C-CSNAP peptides displace CSNAP from the CSN complex

To define whether C-CSNAP can displace the endogenous CSN subunit from the complex, we transiently expressed several versions of C-CSNAP peptides with differing lengths, fused to the fluorescent protein cerulean (Cer) in WT HAP1 cells and cells lacking CSNAP (ΔCSNAP cells) (Fig. 2A). We then performed co-immunoprecipitation analyses, using an antibody against GFP that recognizes the Cer tag (Figure 2B-C). The results indicated that in WT cells, C-CSNAP is pulled down, together with CSN5, indicating its integration within the complex. Moreover, in ΔCSNAP cells all C-CSNAP derivatives were efficiently incorporated into CSN, while in WT cells differential integration was observed. The 16 amino acid fragment of C-CSNAP-Cer displayed the highest degree of association with the CSN complex. Surprisingly, this peptide outperformed the full-length CSNAP-Cer protein in its efficiency in displacing the endogenous CSNAP subunit. Truncations of the 16 amino acid stretch from either end decreased or abolished the incorporation efficiency (Fig. 2B and C).

**Figure 2.**
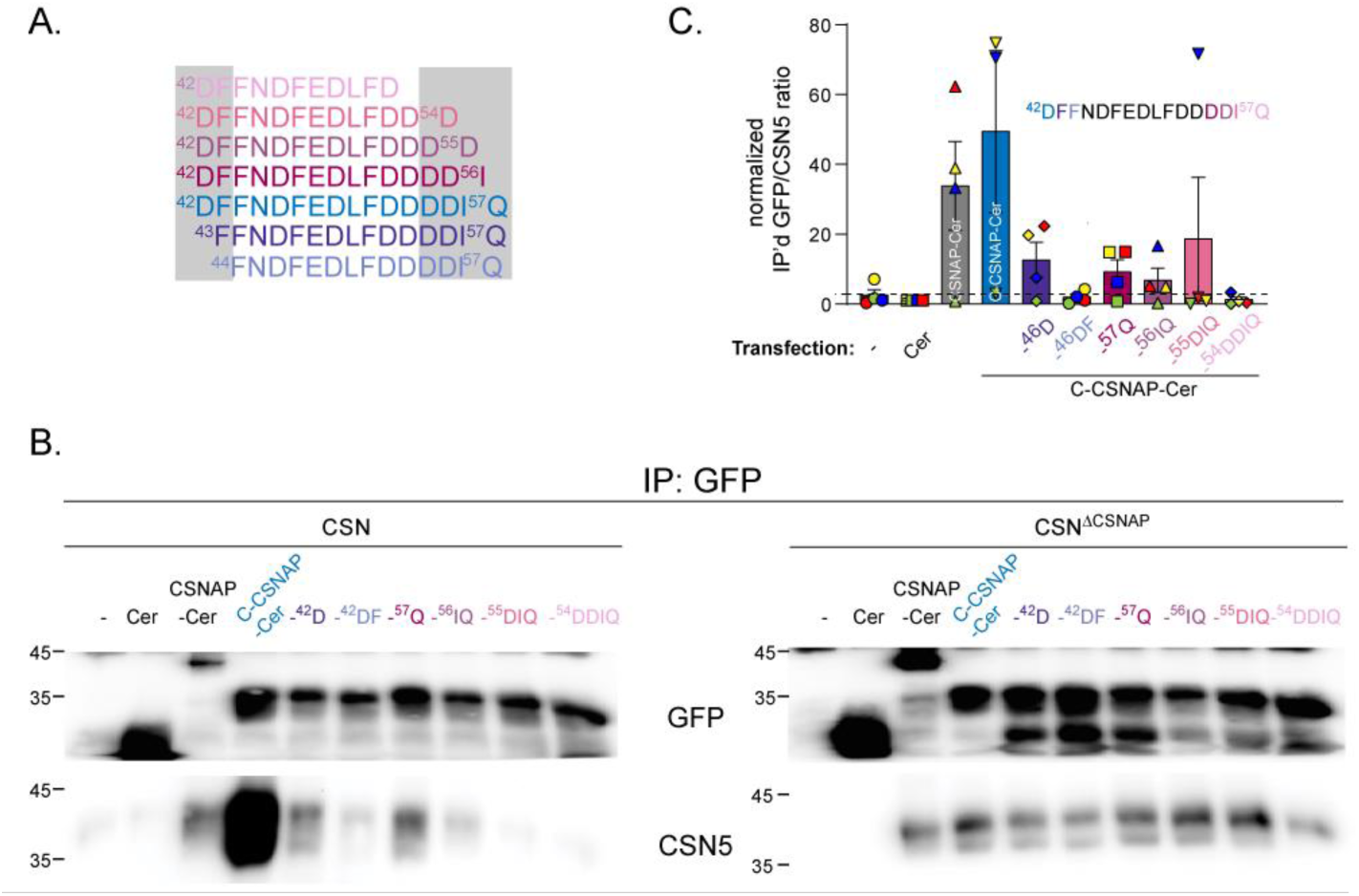
The last 16 amino acids of CSNAP efficiently displace the endogenous CSNAP subunit. A) The different C-terminal sequences that were examined. The 16 amino acid sequence is highlighted in blue, and the N-or C-terminally truncated forms are labelled in tones of purple and pink, respectively. (B) Immuno-precipitation experiments of cells transiently expressing full length CSNAP or truncated versions fused to GFP (cerulean). The last 16 amino acids of C-CSNAP fused to GFP preferentially incorporate into the CSN complex. The efficiency of incorporation of the 16 amino acid C-CSNAP outperforms even that of the full length CSNAP subunit. C) Quantification of the densitometry results of at least three independent immuno-precipitation experiments using an anti-GFP antibody, represented as ratios of immuno-precipitated GFP/ CSN5, followed by normalization to the sample transfected with cerulean, plotted as mean ± standard error of the mean (SEM).

### Optimization of the C-CSNAP peptide length and sequence

To generate an optimized C-CSNAP peptide we used the peptide array methodology. The array was designed to screen for the optimal peptide length, sequence and stability (Fig. 3A). A recombinant CSN complex was expressed, purified and screened for binding to the peptide array by using an anti-CSN3 antibody. To rule out non-specific binding of the probing antibody, a control experiment was carried out without the addition of the CSN complex (Fig. 3A right panel). The anti-CSN3 antibody bound to one spot, which was disregarded in further analysis. The intensity of each spot, reflecting the interaction strength, was normalized to the 16 amino acid C-terminal fragment of CSNAP (Fig 3B-G).

**Figure 3.**
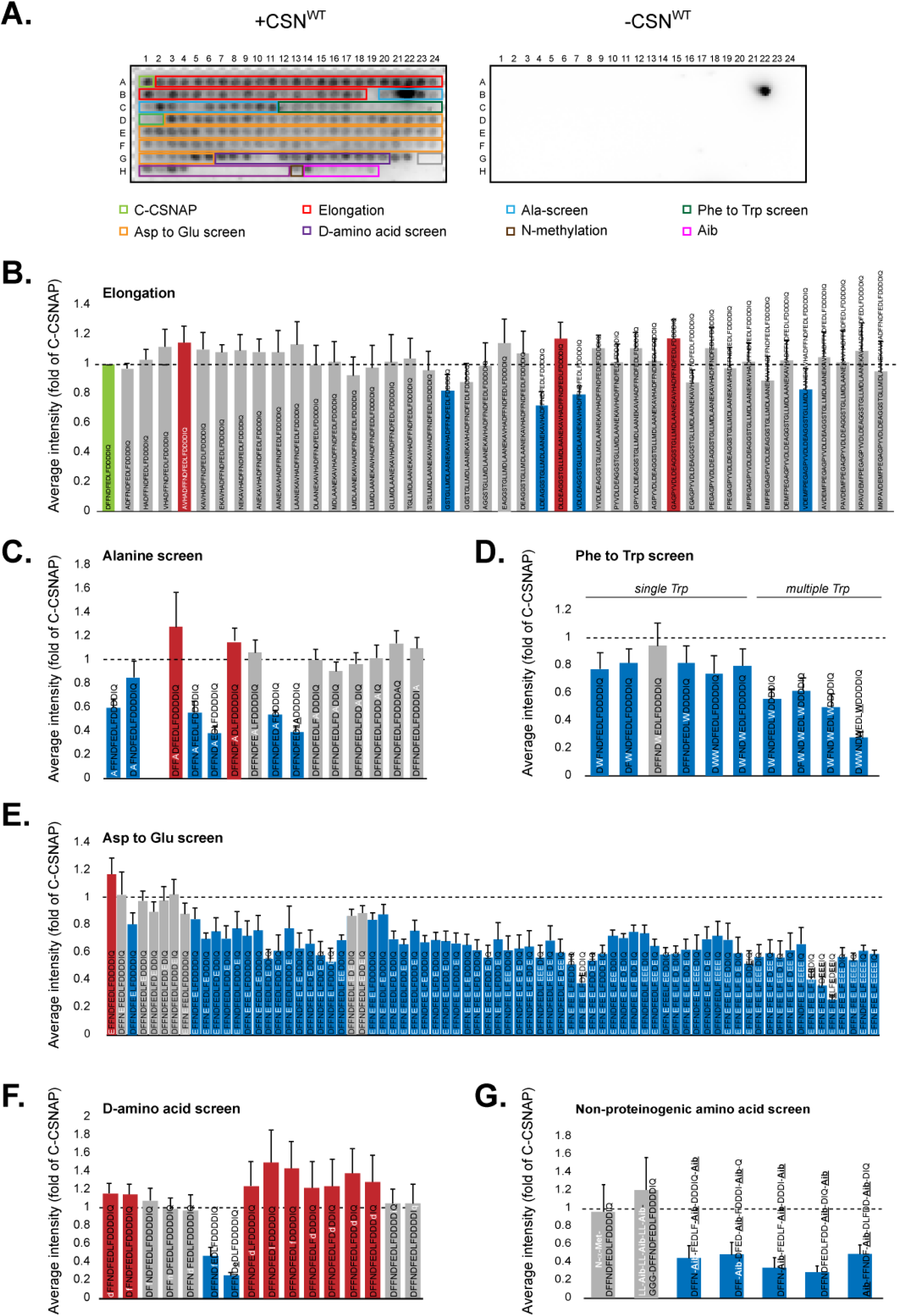
Peptide array screen of modified C-CSNAP peptides. C-CSNAP peptide and its derivatives were synthesized on a microarray. The array was blocked and incubated with or without the CSN complex and probed using an anti-CSN3 antibody and HRP-conjugated secondary antibody. To assess non-specific binding of the antibodies to the array, anti-CSN3 and anti-GAR-HRP were used without the addition of the CSN. Signal intensity of each spot was measured and normalized to the intensity of the spot of C-CSNAP (A1). Measured values for each spot were averaged from 6 arrays, and plotted with standard errors of the mean (SEM). Red bars indicate increased binding compared to C-CSNAP (> 1.14-fold, average - SEM>1), and blue bars represent weaker interaction (<-1.14-fold, average + SEM<1). In (C) results for spot B22 was disregarded, due to non-specific CSN binding to this spot. (A) A representative image of the peptide array (left) and the background (right). (B) Elongation of the C-terminal 16 amino acids (green) with one residue at a time based on the full length sequence. The data highlights stronger binding to CSN when the sequence is 20 amino acids long. (C) The bar plot shows the key positions in which single residue substitution with alanine significantly reduced the binding to CSN. (D) Phe substitution screen to Trp demonstrates the specificity of the Phe residues in CSN binding, the more Phe are substituted with Trp the interaction decreased. (E) Asp to Glu substitution emphasizes the importance of Asp residues. Single substitution increased slightly the strength of binding to CSN, while multiple substitutions significantly weakened interaction is most cases. (F) Single D-amino acid substitution at various positions. Substitution of Phe47 significantly reduced binding to CSN, while replacement of Leu50 to Asp55 promoted interaction. (G) Incorporation of non-proteinogenic amino acids disrupted the CSN-C-CSNAP interaction.

Sequential elongation of the C-CSNAP peptide towards the N-terminal enhanced the binding to CSN, with the addition of four amino acids showing the highest signal (Fig. 3B). This is likely due to the polar interaction that is formed in this peptide between ^CSNAP^His41 and ^CSN3^Ser127, as revealed by the predicted structure (Fig. 1). Furthermore, alanine scanning, in which each residue in C-CSNAP was substituted to alanine, was performed in order to determine the residue specific contribution to the CSN interaction interface. The assay showed that substitution of the negatively charged (Asp42 and Asp46) and hydrophobic residues (Phe47, Leu50 and Phe51) reduced the binding to the complex. These results correlate with the model (Fig. 1B-D), as the aspartic acid residues are solvent exposed and hence disfavor replacement to a non-polar residue, while the bulky hydrophobic residues are buried within the CSN3 interface and their substitution to alanine forms cavities. We also examined the impact of conservative substitutions by replacing the aromatic residue phenylalanine to tryptophan and the negatively charged aspartic acid residue to glutamic acid, in attempts to improve binding (Fig. 3D and E). Decreased tolerance to substitution was detected upon replacement of phenylalanine to the bulkier and less hydrophobic tryptophan residue, supporting the notion of tight packing of the four tryptophans in C-CSNAP (Fig. 1D). The exchange of Asp to Glu at position 42 (D42E), however, increased the binding suggesting that the larger solvent exposed increase polar surface of glutamic acid eases the formation of hydrogen bonds with the aqueous solvent (Fig. 1E).

Next, to increase the analyzed chemical space, we examined the ability of peptides bearing non-canonical amino acids. i.e. D-amino acids and the helix-promoting *α*-aminoisobutyric acid (Aib), to bind CSN. Peptide sequences containing D-amino acids are attractive for our use as they are highly resistant to degradation by proteases^30,31^. However, the impact of such replacement may interfere with protein interaction, and hence we examined the effect of a single L-amino acid replacement to D-amino acid on the binding capacity to CSN. The results pointed to the importance of Phe47 and Glu48 in CSNAP-CSN interaction, as their substitution reduced significantly the binding of the peptides (Fig. 3F). On the other hand, D-amino acid substitution of Leu50 to Asp55 exhibited enhanced CSN binding, by more than 20%, in comparison to the original peptide. Single residue D-amino acid substitutions are known to disrupt *α*-helical structure^30^ and given the intrinsically unstructured conformation of C-CSNAP (Fig. 1F), such replacements are probably stabilizing the unstructured conformation of this region, suggesting that such partial D-amino acid substitution may be useful for *in-vivo* activity and stability. Incorporation of an Aib residue, in which the α-hydrogen atom of alanine is replaced with a methyl substituent is commonly used for promoting helicity^32^. This substitution did not significantly affect the ability to interact with CSN when introduced at the first two positions of the peptide, but attenuated binding in all internal substitutions. This observation is in agreement with the impact of the D-amino acid substitutions, highlighting the prevalence of an unstructured conformation of C-CSNAP. Lastly, N-terminal methylation of the peptide, which improves pharmacokinetic properties^33^, did not affect the interaction with CSN.

The same array was also reacted with the CSN^ΔCSNAP^ complex (Supplementary Fig. 2). The data indicate that the interaction of CSN with C-CSNAP peptides is more stringent than CSN^ΔCSNAP^, as expected from the requirement to expel the endogenous CSNAP subunit from the complex prior to peptide binding. Taken together, the peptide array analysis not only provided insight into the optimal C-CSNAP peptide characteristics, it also validated and supported the predicted model.

### C-CSNAP peptides substitute the endogenous subunits

We next focused on exploring the ability of C-CSNAP peptides to displace CSNAP. To this end, based on the peptide array results we selected four peptides, the 16 residue C-terminal fragment (^42^DFFNDFEDLFDDDDIQ^57^, 16AA C-CSNAP), and three of the peptides that exhibited higher binding affinities: the 20 amino acid peptide (^38^AVHADFFNDFEDLFDDDDIQ^57^, 20AA C-CSNAP), a peptide containing the substitution of Asp to Glu at position 42 (D42E C-CSNAP), and a peptide in which Leu50 was exchanged with a D-amino acid (dLeu50 C-CSNAP) (Fig. 4). We incubated the CSN complex with each one of the peptides, bound the complex to streptactin beads (StrepIIx-CSN3) and removed the unbound CSNAP fraction by washing. Next, the complex was separated into its constituent subunits using a monolithic column^34^. The eluted subunits were sprayed directly into the mass spectrometer for intact protein mass measurements. As a negative control, we used the CSN complex without the addition of C-CSNAP. Analysis of the spectra validated the presence of charge series corresponding in mass to the four C-CSNAP peptides, along with the other CSN subunits, indicating that all the designed peptides were able to displace the endogenous CSNAP subunit (Fig. 4, Supplementary Fig. 3). However, the 20AA C-CSNAP peptide displayed a higher capacity for dislocating CSNAP, as shown by its increased abundance relative to the other peptides (Fig. 4C). The incorporation of 20AA C-CSNAP within the CSN complex was also validated by native-MS/MS analysis (Fig. 4D).

**Figure 4.**
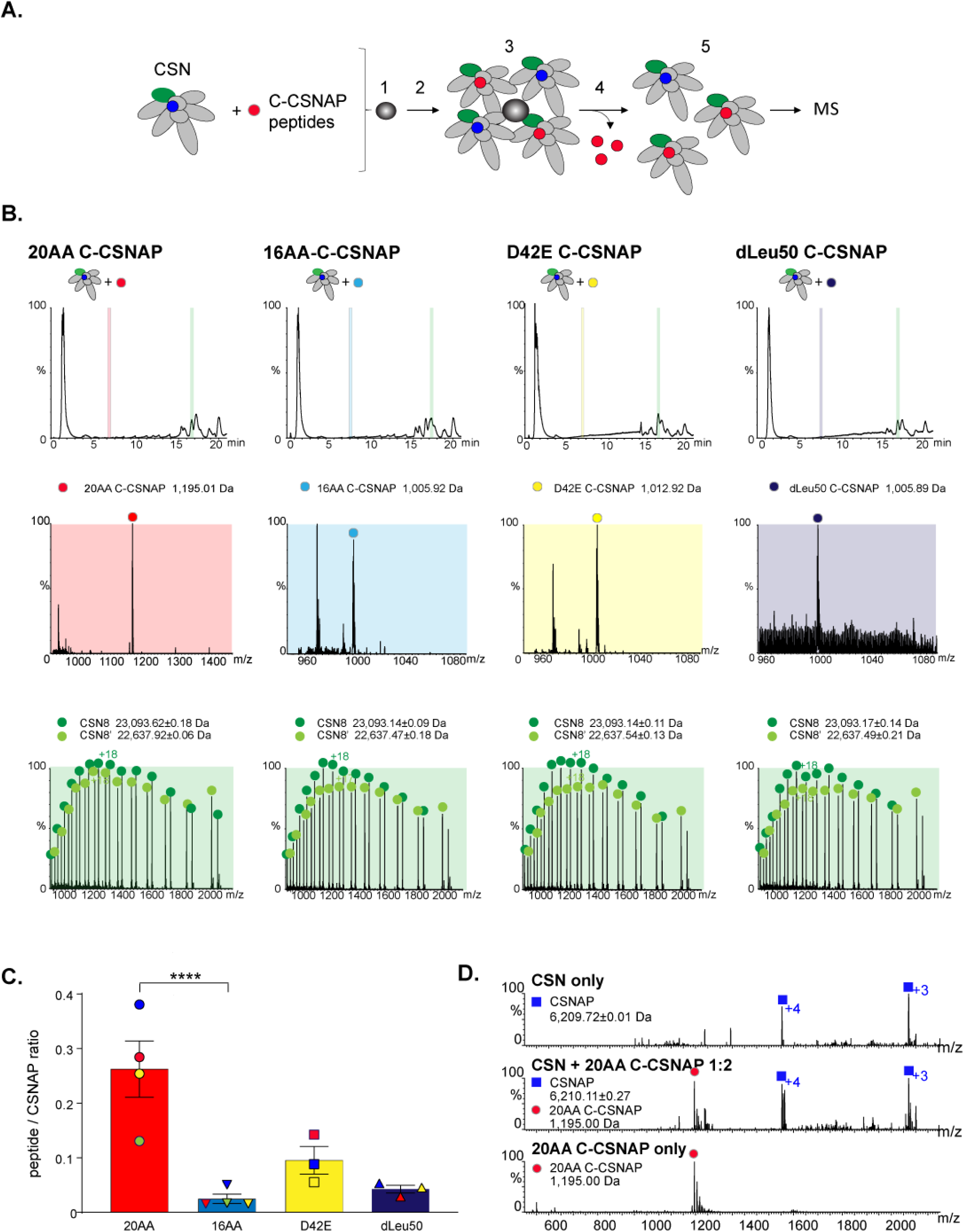
C-CSNAP peptides displace the full length subunit. (A) Graphic scheme of the displacement experiments using C-CSNAP peptide variants and recombinant CSN. CSN was incubated with 2-fold molar excess of 16AA C-CSNAP or modified peptides prior to coupling to streptactin beads (1) for 2.5 hours (2). Streptactin-bound CSN (3) was washed three times (4) to remove free peptides, and CSN was eluted from the beads (5) for MS analysis. (B) In this analysis the CSN was separated into its component subunits, using a monolithic column under denaturing conditions. The eluted subunits are directed straight into the mass spectrometer for intact protein mass measurements. (B) Masses corresponding to all four peptides were detected (highlighted with red, light blue, yellow and dark purple for 20AA, 16AA, D42E and dLeu50 C-CSNAP, respectively). Representative spectra of the canonical CSN subunit, CSN8 (green). The two different series of peaks in the ESI-MS spectrum, correspond to full-length CSN8 and its alternative translation initiation site isoform^34^. (C) The bar plot shows average ratios of intensities corresponding to masses of each peptide and CSNAP after incubation with CSN with SEM. Significance was calculated from a minimum of 3 experiments for each peptide using 1-way ANOVA, yielding a value of p<0.005, followed by Dunnett’s multiple comparisons test ****p<0.001. (D) The CSN complex was incubated either with or without the 20AA C-CSNAP peptide at a ratio of 1:2 and then analyzed by tandem MS. MSMS analysis of the intact CSN complex resulted in dissociation of CSNAP (upper panel, dark blue squares). Activation of the CSN complex following preincubation with 20AA C-CSNAP resulted in dissociation of both CSNAP and 20AA C-CSNAP (middle panel, dark blue squares and red circles, respectively), demonstrating that prior to MSMS analysis, 20AA C-CSNAP was physically associated with the CSN complex. As a control, C-CSNAP was measured alone, and was detected as a doubly charged ion (lower panel, red circle).

### C-CSNAP peptides reduce cellular proliferation

To clarify the ability of C-CSNAP peptides to displace CSNAP in cells, we stably expressed two versions of CSNAP, the 16AA and 20AA C-CSNAP peptides, fused to the fluorescent protein Cer in HAP1 cells. We then performed reciprocal co-immunoprecipitation analyses, using antibodies against CSN3 and GFP (Fig. 5A and Supplementary Fig. 4). The results indicated that both C-CSNAP peptides are integrated efficiently into the CSN complex. However, based on band intensities the shorter peptide, 16AA C-CSNAP, displayed a higher ability to integrate into CSN, compared to the 20AA peptide. This is reflected by the higher intensity of the GFP band upon CSN3 pull-down, and by the more intense CSN3 band following GFP co-immunoprecipitation.

**Figure 5.**
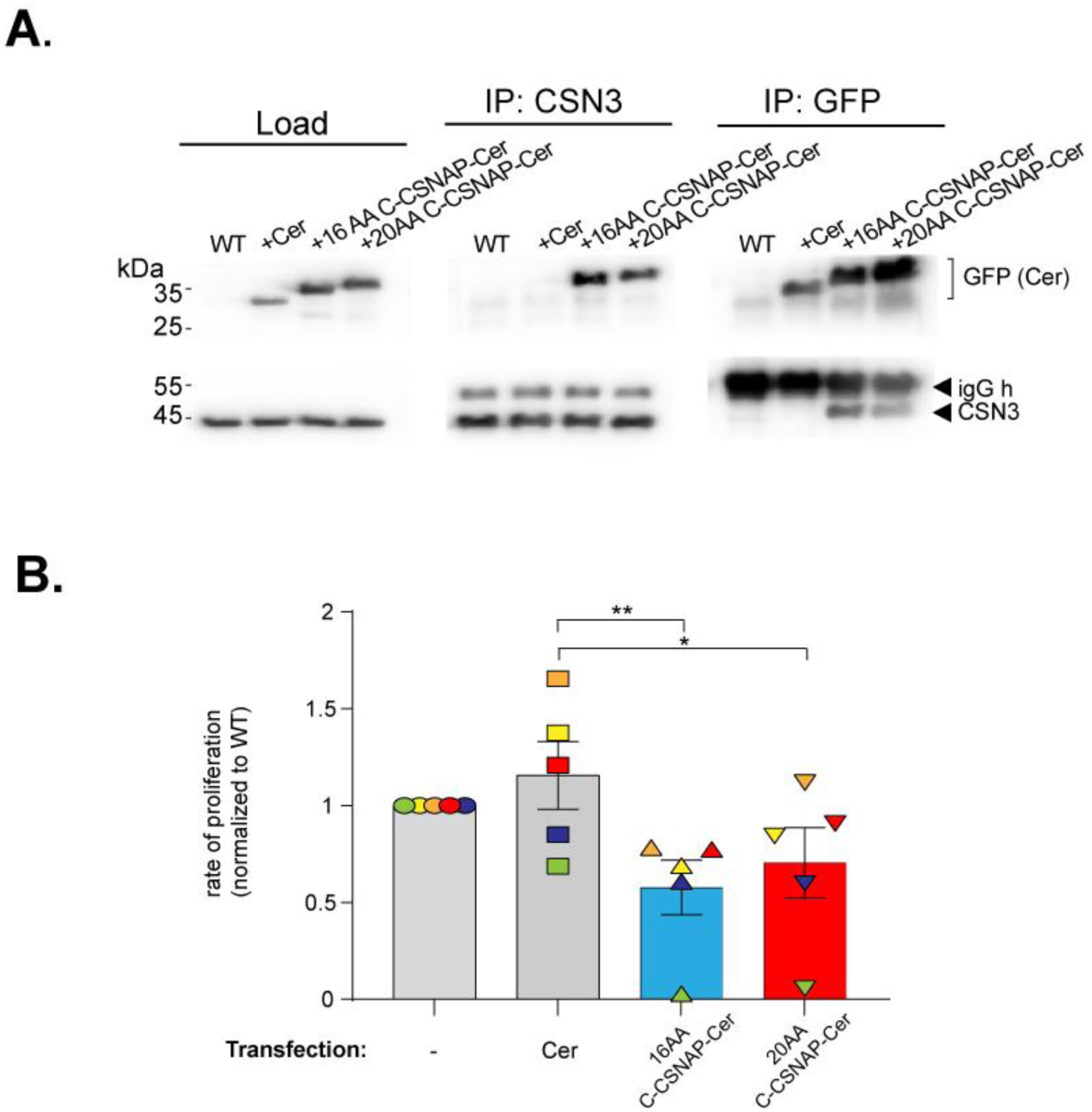
Expression of C-CSNAP reduces cellular proliferation rate. (A) The CSN complex was immuno-precipitated using anti-GFP antibody recognizing cerulean (Cer) from lysates of cells stably expressing the cerulean-fused 16AA and 20AA C-CSNAP in HAP1 WT cells, but not when Cer is expressed alone. Similarly, reciprocal immuno-precipitation by an anti-CSN3 antibody pulled-down only the cerulean-fused 16AA or 20AA C-CSNAP. (B) Stable overexpression of 16AA and 20AA C-CSNAP-cerulean in HAP1 WT cells reduces their rate of proliferation. The bar plot shows average proliferation rates of each cell line using 5 independent replicates with SEM. One-way ANOVA with Dunnett’s multiple comparisons test was used to compare means *p<0.05, **p<0.01.

Previously we have shown that ΔCSNAP cells proliferate at a slower rate compared to WT cells^23,24^. We hypothesized, that displacement of the endogenous CSNAP by the exogenously expressed C-CSNAP peptide fused to cerulean, will induce a similar effect, i.e. reducing cell proliferation, similar to ΔCSNAP cells. We, therefore, compared the proliferation rates of WT, ΔCSNAP, 16AA and 20AA C-CSNAP-Cer expressing cell lines in resazurin-based viability assays. The data show that there are significant reductions in cell proliferation in both 16AA and 20AA C-CSNAP clones compared to the WT cells or cells expressing full-length CSNAP fused to Cer cells (Fig. 5A). In summary, this result confirmed the cellular ability of a C-CSNAP derived peptide to dislocate the endogenous subunit and thus attenuate CSN activity.

## Discussion

Here we showed that a C-terminal peptide of CSNAP can displace the endogenous CSNAP subunit. We demonstrated that the peptide is incorporated within the complex, and that upon its binding cell proliferation is reduced, thereby, mimicking the phenotype observed upon deletion of CSNAP^24^. The results are based on an AlphaFold2 generated high confidence prediction model of CSN1, CSN3, CSN8 and CSNAP that agrees very well with our mutational analysis^24^ and peptide array results (current study) as well as with the previously solved X-ray CSN^ΔCSNAP^ structure^27^ and chemical cross-linking constraints^26^. The model provides important insight into the structural details of CSN-bound CSNAP. Coupling the structural and experimental results revealed that a 16 amino acid residue peptide displays the highest cellular capacity for substituting the full length subunit from the CSN. Overall, our results suggest that inhibition of the CSN complex can be achieved by a peptide based approach.

In recent years, pharmaceutical research is revisiting the use of peptides as therapeutics, especially for inhibiting protein-protein interactions that are more difficult to therapeutically target than the interactions of globular proteins with other biomolecules^35-37^. This is due to the fact that peptide based drugs benefit from high target specificity, strong binding affinity, low immunogenicity, a lower potential for drug-drug interaction, and high tolerability and safety, whilst effective therapeutic outcomes can be reached with only a small concentration of peptide^38,39^. Moreover, advances in structural biology, recombinant biologics, and new synthetic and analytic technologies have significantly accelerated peptide drug development^40^. There are still major challenges to overcome before wide clinical application of peptides can be considered. Susceptibility to proteolytic degradation, delivery and rapid renal clearance are known limitations of peptide-based therapies^41^. Despite these challenges, more than 80 therapeutic peptides have reached the global market to date, and over 170 peptides are in active clinical development, with many more in preclinical studies^40^, emphasizing the prospective our approach.

IDPs often fold into ordered states upon binding to their physiological interaction partners^42^. The C-terminal region of CSNAP, however, remains disordered in its CSN-bound state, as predicted by the model and verified by experimental input. Although less common, multiple examples of IDPs remaining disordered upon binding exist^43^. For instance, the binding of the intrinsically disordered regulatory region of CFTR to NBD1^44^, the binding of the IDP Sic to Cdc4^45^, or the flexible stretch that remains in the structure of p21 and p27 when bound to CDK^46,47^. Even though C-CSNAP does not adopt a secondary structure, it seems to be tightly anchored to CSN3 and CSN8 by its 5 aromatic residues that anchor to CSN3 and the complementary electrostatic interactions. Thus, it likely adopts a confined bound conformation rather than populating an ensemble of conformational states. The use of peptides for targeting protein-protein interactions mediated by IDPs is still in its infancy, however, examples of this strategy are already available, such as the interaction of *α*-synuclein and TPPP/p25^48^, AF4 and AF9^49^, p53 to MdM2^50^ as well as the interactions of the histones H4 and H2A^51^, and NURP1^52^. Such peptides can be utilized as starting structures for developing peptidomimetics. Methodologies relevant for IDP peptidomimetics are already being developed possessing protease resistance and increased cell permeability^53^, supporting their potential use of this approach for pharmaceutical intervention.

In summary, the main finding of this study is the demonstration that C-CSNAP peptides inhibit CSN activity. The inhibition of CSN is an attractive goal for cancer research since this complex is associated with cancer development and progression. The role of CSN in tumorigenesis is likely indirect and is linked to coordinating CRLs function. Cell division and signal transduction are tightly regulated by CRL activity, thus inhibiting CRLs through down regulation of CSN is a promising therapeutic avenue. The impact of CSN inhibition on protein degradation is more restricted compared to the proteasome complex, as only the degradation of CRLs substrates is attenuated, possibly leading to reduced resistance and side effects compared to the treatment of proteasome inhibitors. Taken together, the peptide based strategy we propose here has the potential to be further developed as lead compounds opening a new avenue for selective CSN inhibition^54,55^.

## Materials and Methods

### AlphaFold model prediction

To implement AlphaFold2^28,29^ locally, we used an adapted code written by ColabFold^56^. The run used the five model parameters, 7 recycle rounds, and no templates or Amber relaxation. Multiple-sequence alignments were generated through the MMseqs2 API server^57-59^.

### Expression of GFP constructs

The C-terminal ^42^DFFNDFEDLFDDDDIQ^57^ (C-CSNAP) sequence of CSNAP was generated in pHyg vector coding for full length CSNAP, fused the N-terminus of cerulean through a short Gly-Ser-Gly-Ser linker, by deleting ^1^M-^41^A residues of CSNAP, using Q5 mutagenesis kit (NEB). N-(-^42^D, -^42^DF) and C-terminal truncations (-^57^Q, -^56^IQ, -^55^DIQ and -^54^DDIQ) were performed on phyg-C-CSNAP-cerulean using the Q5 mutagenesis kit (NEB). HAP1 WT cells were transfected with 5 *μ*g plasmid DNA using JetPRIME (Polyplus), and culture medium was changed 4 hours post-transfection. Stably expressing WT cell lines were prepared by applying hygromycin (0.4 mg/ml) selection for 7 days following transfection, then the pool of surviving cells were sorted on a FACSAriaIII cell sorter for cerulean positive cells.

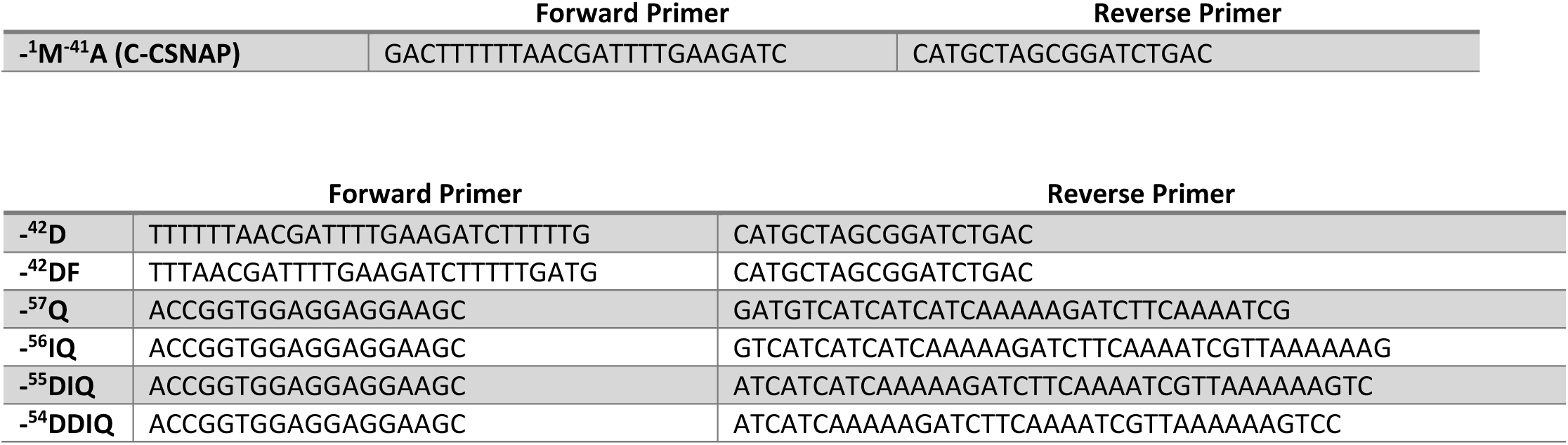

Expression levels were monitored by loading 30*μ*g of each lysate on 12% Laemmli SDS-PAGE, transferred to PVDF membrane and probed with anti-GFP antibody (Abcam -ab290).

### Immunoprecipitation (GFP)

HAP1 WT and ΔCSNAP cells were transiently transfected for 72 hours with pHyg-C-CSNAP-cerulean, N- or C-terminally truncated forms, and cells were lysed (50mM Tris-HCl pH 7.4, 150 mM NaCl, 0.5% NP-40, 2.5 mM Na-pyrophosphate, 1 mM *β*-glycero-phosphate and 1mM Na-ortho-vanadate supplemented with PMSF, benzamidine and pepstatin). For each immuno-precipitation 250 *μ*g lysate was incubated with 1*μ*l anti-GFP (Abcam -ab290) in 500 *μ*l (TBS) overnight, then with 20 *μ*l Protein-G Sepharose for 1 hour, and bound proteins were eluted with 40 *μ*l 2x Laemmli sample buffer.

### Mass spectrometry

The monolithic-LC-MS approach applied on the recombinant CSN complex was performed as previously described^34^. Briefly, 76.5 pmoles of recombinant CSN^WT^ was incubated with 2-fold molar excess (75 pmoles) of 16AA, 20AA, dLeu50 or D42E C-CSNAP peptides in 50 *μ*l CSN wash buffer (50 mM Tris-HCl pH 7.4, 250 mM NaCl, 2 mM EDTA, 2 mM DTT) 3 hours at 37°C. CSN^WT^ alone or with each peptide, and peptides alone were incubated with 10 *μ*l Streptactin beads (IBA) in 500 *μ*l at 4°C for 2.5 hours. The beads were washed with 200 *μ*l CSN wash buffer, then twice with 200 *μ*l 10mM HEPES pH7.5, and bound CSN was eluted with 30 *μ*l 2.5mM destiobiotin in 10mM HEPES pH7.5 for 30’ at 37C. 5 *μ*l of the 30 *μ*l was injected to the monolithic column and denatured with a gradient 10-50% acetonitrile +0.035% TFA + 0.05% formic acid (20 min). Ratio between CSNAP (sum of intensities of charge states +5, +4, +3) and bound peptide (+2) was calculated for each sample in 3 or 4 replicates, and statistical significance between CSN-bound CSNAP/peptide ratios were compared using Student’s t-test (unequal variance), p<0.05).

### Recombinant CSN expression

Recombinant CSN^ΔCSNAP^ and CSN was expressed in Sf9 insect cells and purified as described in ^60^.

### Peptide array

Optimization of C-CSNAP sequence for enhanced binding to recombinant (r)CSN^ΔCSNAP^ was screened by printing the modified sequences of CSNAP (6xHis-tag or sequentially truncated) or C-CSNAP (Phe to Trp, Asp to Glu, Ala screen, D-amino acid screen, N-*α*-methylation, Tat-, Arg7, and 6xHis-tag, aminoisobutyric acid (Aib) on INTAVIS Celluspots array (2×384spots). The array was blocked with 5% skim milk in TBS-0.5% Tween 20 (TBS-T) for 4 hours at room temperature, washed 3 times in TBS-T, then incubated with 0.5 *μ*M CSN^ΔCSNAP^ in 5% milk-TBS-T. Bound CSN^ΔCSNAP^ was visualized by incubating with anti-CSN3 (ab79698) antibody (1:10,000, overnight) and goat-anti-rabbit-HRP secondary antibody (1:10,000, 1 hr).

### Cell proliferation assays

HAP1 WT cells stably expressing cerulean, 16AA and 20AA C-CSNAP-cerulean were trypsinized and counted. Cells from each cell group were plated in a 24 well plate (4 replicates, 5000 cells/well) for proliferation assay. Resazurin based assays were performed as described in ^23^, 24 hours following transfection. Fluorescence intensities at 560/600nm (ex/em) were measured, and average intensities of 4 replicates in each plate were calculated, and normalized against WT after background subtraction, then normalized values from each replicate experiments were compared on a bar graph with SEM. Significance was calculated using paired Student’s t-test (*p<0.05).

## Acknowledgments

MS is grateful for the support of the Sagol Institute for Longevity Research grant and a Moross Proof-of-Concept grant. MS is the incumbent of the Aharon and Ephraim Katzir Memorial Professorial Chair. SJF was supported by the Dr. Barry Sherman Institute for Medicinal Chemistry.

## Supplementary Figures

**Supplementary Figure 1.**
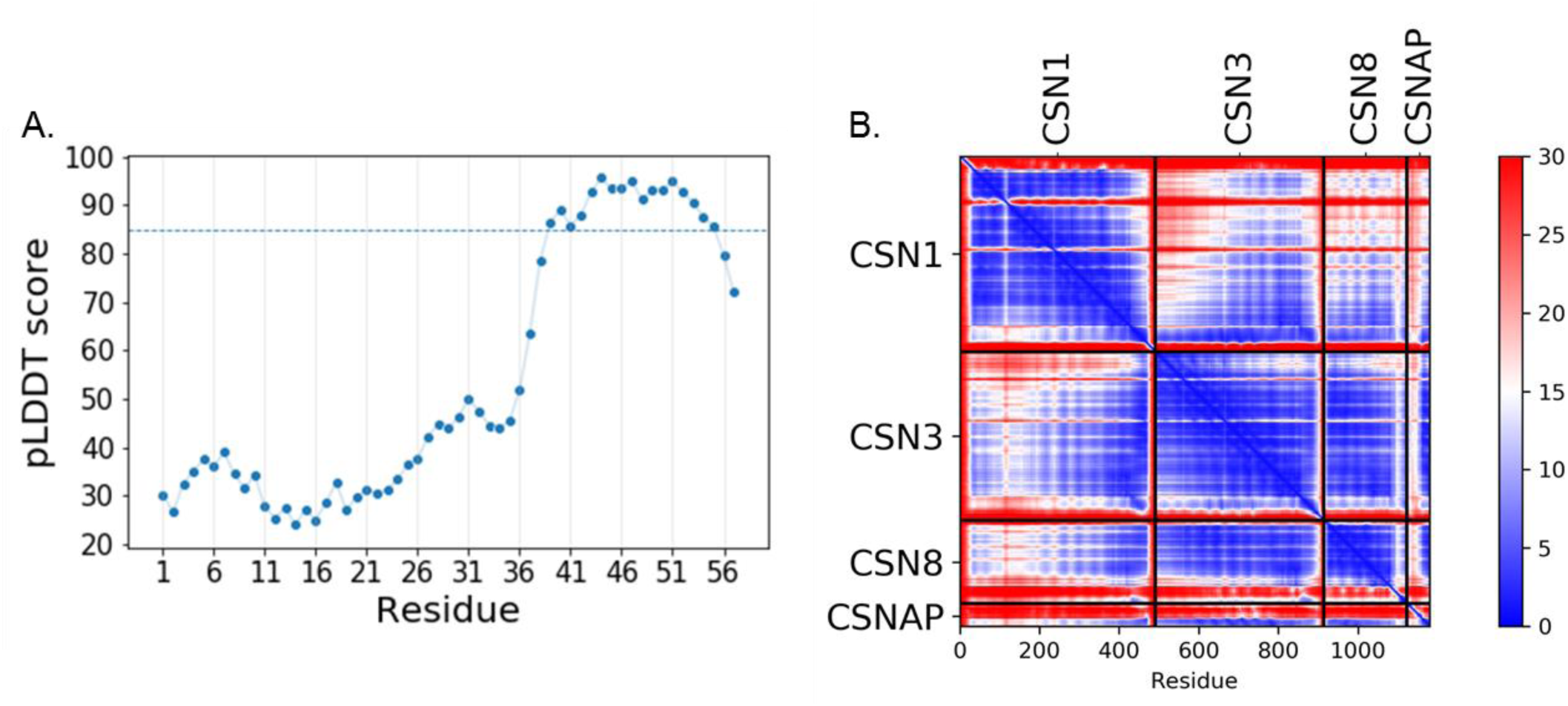
The C-terminal CSNAP structure displays high AlphaFold confidence parameters. (A) pLDDT scores for CSNAP. Regions with pLDDT > 85 are expected to be modeled to high accuracy which is the case with the C-terminus tail of CSNAP. (B) PAE scores for the CSN1, CSN3, CSN8, CSNAP combined complex. When PAE is high (red) the relative positions of the residues in the 3D structure is uncertain. The C-terminus domain of CSNAP shows high certainty scores (blue) in the interface of the CSN3 and CSN8 domains.

**Supplementary Figure 2.**
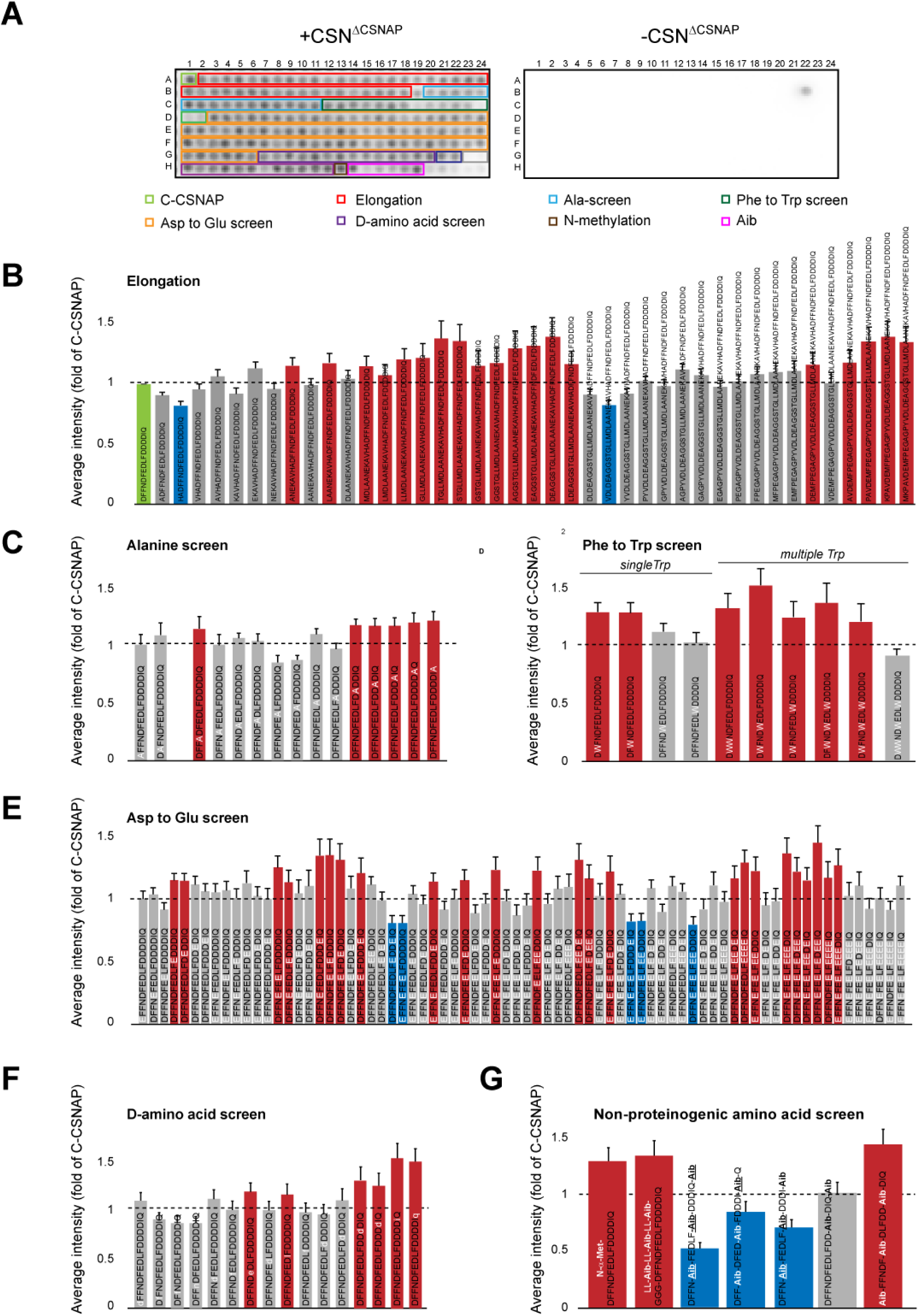
Peptide array screen of modified C-CSNAP peptides. C-CSNAP peptide and its derivatives were printed on a on a microarray. The array was blocked and incubated with or without the CSN^ΔCSNAP^ complex and probed using an anti-CSN3 antibody and HRP-conjugated secondary antibody. To assess non-specific binding of the antibodies to the array, anti-CSN3 and anti-GAR-HRP were used without the addition of the CSN. Signal intensity of each spot was measured and normalized to the intensity of the spot of C-CSNAP (A1). Measured values for each spot were averaged from 6 arrays, and plotted with standard errors SEM. Red bars indicate increased binding compared to C-CSNAP (> 1.15-fold, average - SEM>1), and blue bars represent weaker interaction (<-1.15-fold, average - SEM<1). In (C) results for spot B22 was disregarded, due to non-specific CSN binding to this spot. (A) A representative image of the peptide array (left) and the background (right). (B) Elongation of the C-terminal 16 amino acids (green) with one residue at a time based on the full length sequence. (C) The bar plot shows the key positions in which single residue substitution with alanine significantly weakens binding to CSN. (D) Phe substitution screen to Trp and (E) Asp to Glu replacements. (F) Single D-amino acid substitution at various positions. (G) Incorporation of non-proteinogenic amino acids.

**Supplementary Figure 3.**
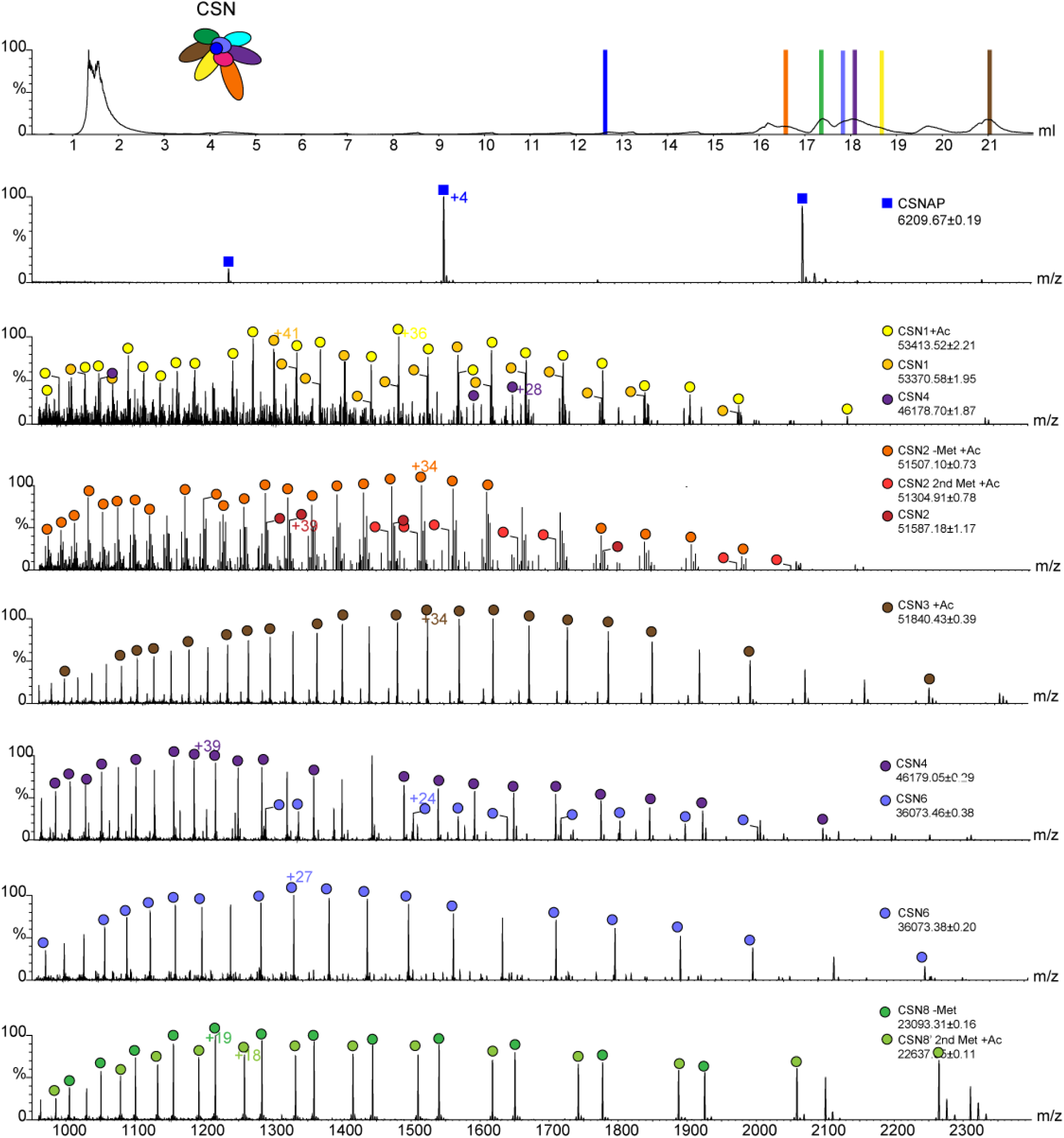
Representative CSN subunits identified in the displacement experiment (Figure 4). The recombinant CSN complex was bound to streptactin beads and after elution separated into its composing subunits, using a monolithic column under denaturing conditions followed by on-line MS analysis. Peaks representing the eluted CSN subunits are highlighted in color in the top panel and correlated with the recorded ESI-MS spectra below.

**Supplementary Figure 4.**
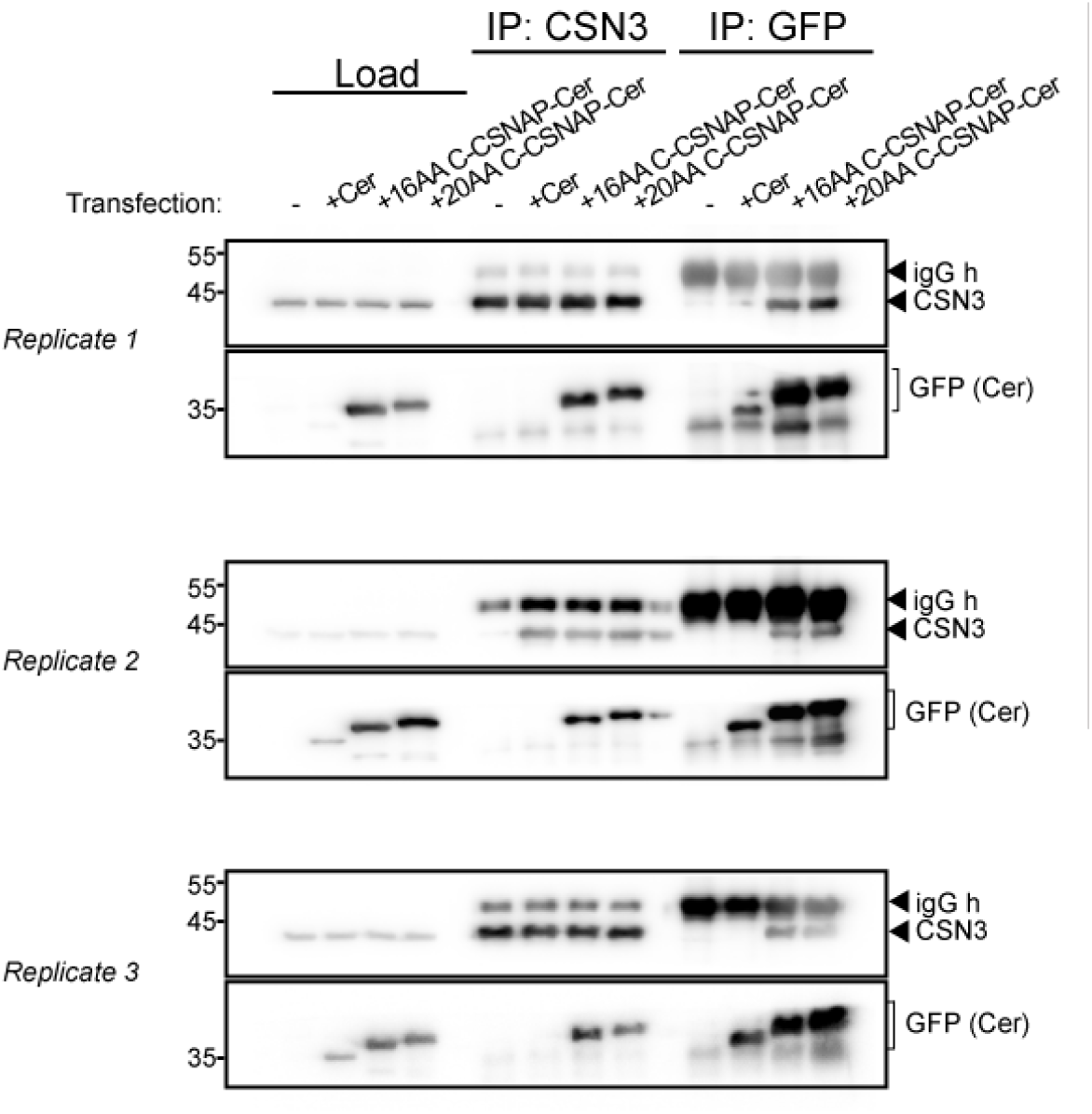
Replicas of reciprocal co-immunoprecipitation experiments, indicate the integration of the C-CSNAP peptides within CSN. The CSN complex was immuno-precipitated using anti-GFP antibody recognizing cerulean (Cer) from lysates of cells stably expressing the cerulean-fused 16AA and 20AA C-CSNAP in HAP1 WT cells, but not when Cer is expressed alone. Similarly, reciprocal immuno-precipitation by an anti-CSN3 antibody pulled-down only the cerulean-fused 16AA or 20AA C-CSNAP.

